# The monitoring system is attuned to others’ actions during dyadic motor interactions

**DOI:** 10.1101/2021.04.01.438029

**Authors:** Quentin Moreau, Gaetano Tieri, Vanessa Era, Salvatore Maria Aglioti, Matteo Candidi

## Abstract

Successful interpersonal motor interactions necessitate the simultaneous monitoring of our own and our partner’s actions. To characterize the dynamics of the action monitoring system for tracking self and other behaviors during dyadic synchronous interactions, we combined EEG recordings and immersive Virtual Reality in two tasks where participants were asked to coordinate their actions with those of a Virtual Partner (VP). The two tasks differed in the features to be monitored: the *Goal* task required participants to predict and monitor the VP’s reaching goal; the *Spatial* task required participants to predict and monitor the VP’s reaching trajectory. In both tasks, the VP performed unexpected movement corrections to which the participant needed to adapt. By comparing the neural activity locked to the detection of unexpected changes in the VP action (other-monitoring) or to the participants’ action-replanning (self-monitoring), we show that during interpersonal interactions the monitoring system is more attuned to others’ than to one’s own actions. Additionally, distinctive neural responses to VP’s unexpected goals and trajectory corrections were found: goal corrections were reflected both in early fronto-central and later posterior neural responses while trajectory deviations from the expected movement were reflected only in later and posterior responses. Since these responses were locked to the partner’s behavior and not to one’s own, our results indicate that during interpersonal interactions the action monitoring system is dedicated to evaluating the partner’s movements. Hence, our results reveal an eminently social role of the monitoring system during motor interactions.

*Significance Statement:* Non-verbal synchronous interpersonal interactions require the monitoring of both our actions and those of our partners. Understanding the neural underpinnings of this ability with a focus on the dynamics between self- and other-monitoring is fundamental to the comprehension of social coordination. By combining EEG and immersive Virtual Reality we demonstrate that the monitoring system is more attuned to others’ actions than to our own. In two tasks, we show that the neural activity associated with unexpected corrections in the goal or the trajectory of an action are locked to the partner’s actions rather than to the participants’ subsequent adaptation. This pattern of results highlights a social mode adopted by the monitoring system to handle motor interactions.

## Main Text

### Introduction

Our brain constantly monitors our own actions and their consequences and compares them to the intended ones. This allows adequate adjustments and corrections when our movements happen to be erroneous. A vital component of this monitoring mechanism is the detection and the processing of errors (1–3).

Electroencephalography (EEG) studies have extensively described specific event-related potentials (ERPs) associated with errors. First, Falkenstein and colleagues (4) highlighted a fronto-central ERP (i.e., the Error-Related Negativity – ERN) recorded between 0-150 ms after participants made a mistake in a Go/NoGo task. They also described a later, more posterior, positive potential (i.e., Pe) peaking around 300 ms after the performance of an incorrect action. The Pe was later decomposed into an early and a late component, with the early Pe associated with the involuntary attention switch to novel or unexpected stimuli, and the late Pe reflecting conscious recognition of an error and decision-making vis-a-vis unexpected stimuli (5–7).

Evidence suggests that error processing follows a hierarchical organization. On the one hand, “low-level” sensorimotor errors are defined as discrepancies between motor commands and action effects as movements continuously unfold. On the other hand, “high-level” errors reflect the actual failure of reaching a goal (8). Interestingly, while “high-level” errors are associated with fronto-central activity and the generation of the ERN, “low-level” errors and corrections are primarily related to activity recorded over posterior sites (e.g., the posterior parietal cortex, 8, 9). Accordingly, the detection and correction of “low-level” errors does not generate an ERN and is instead highlighted by the modulation of late parietal potentials such as late Pe (9). In the time-frequency domain, theta (4-7Hz) synchronizations over midfrontal electrodes have been reported during error commission (10–13). More generally, midfrontal theta is considered a marker of “high-level” processes of prediction errors and as a mechanism of cognitive control and behavioral adjustment (14), as confirmed by the use of transcranial alternating current stimulation used to modulate frontal theta activity during post-error adjustment (15).

Notably, the mere observation of someone else performing an error generates time and time-frequency EEG signals that are reminiscent of those detected when performing an error in the first person (16–22). Studies have also shown that observed errors are reflected in the activity of the observer’s motor system (23, 24) hinting at the idea that predictive action simulation supports the ability to detect an error in the actions of others (25, 26). Altogether, these findings suggest a common coding for performed and observed errors, leading to the question of how the monitoring system handles synchronous interpersonal interactions, when both self and others’ behaviors need to be monitored simultaneously.

Motor interactions, ranging from shaking hands to dancing together are omnipresent in our everyday social life and require constant monitoring of our own movements as well as the movements of others (27–29). However, current knowledge regarding the brain dynamics of self-and other- action monitoring in ecological scenarios, where the two mechanisms co-occur, is somehow sparse. The neurophysiological and behavioral correlates associated with errors have only recently been explored in interactive scenarios (30, 31). In the present study, we aim to target action monitoring mechanisms in interactive paradigms with a Virtual Partner (VP). We developed two interactive tasks in a fully immersive VR environment where the participant controlled a virtual body in 1st-person perspective (1PP) while interacting with a VP seen in third-person perspective (3PP). The overall objective of both tasks consisted in controlling the participants’ virtual right upper limb/hand and making it synchronous in pressing a button with the virtual partner in order to turn a lightbulb green. Crucially, in 30% of the trials, the VP performed an unexpected correction of its movement, requiring the participants to quickly react and adapt their action (*Correction factor*). The two tasks differ in the type of corrections performed by the VP. In the *Goal* task, the VP changed the finger used to press one of two different buttons (from using the index finger to using the middle finger or vice-versa, see Figure 1A and Supplementary Video), while in the *Spatial* task, the VP changed its reaching trajectory (from reaching a left button to a right button or vice-versa, see Figure 1B and Supplementary Video). In addition, depending on the block’s instruction, participants were asked to perform the task in an imitative or in a complementary way (*Interaction-type factor*, see Figure 1 and Supplementary Video).

**Figure 1.**
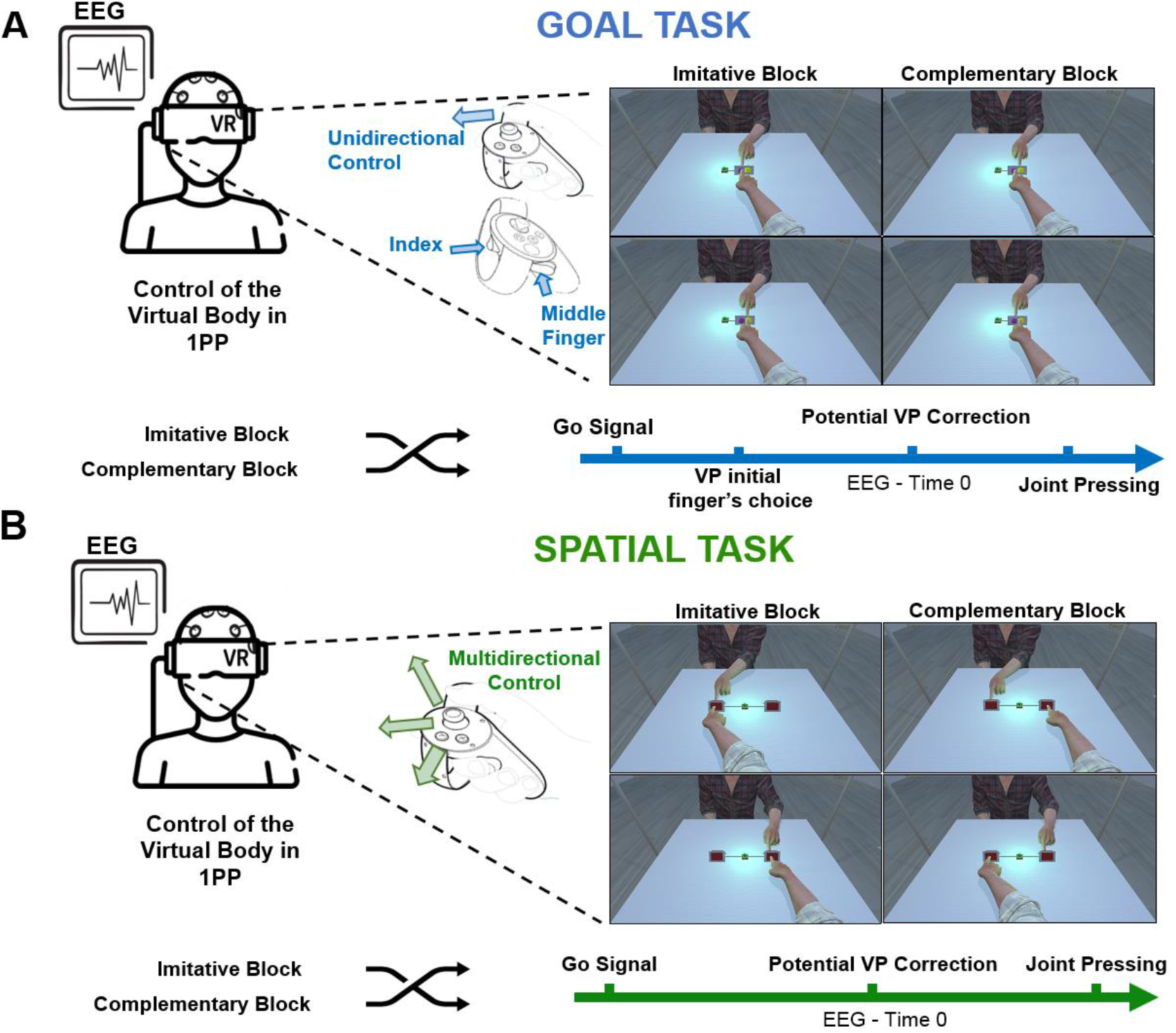
Experimental tasks. In both tasks, participants were put into a fully immersive virtual environment and could move their virtual right arm and hand (first person perspective – 1PP) by means of a controller to interact with a Virtual Partner (VP). The general aim for both tasks was to press a target button simultaneously with the VP. **A)** In the Goal task, participants used the analogic stick of the controller with the thumb to move the virtual hand forward and, by pressing the index and middle trigger button of the right Oculus controller, they could raise either their virtual index or their virtual middle finger. Participants were asked to reach and press the same-colored button using the same finger as virtual partner (Imitative Block) or to press a different colored button, using the opposite finger with respect to the VP (Complementary Block). Along the trial, the VP would initially raise a finger (VP initial finger’s choice), and in 30% of the trials, correct its initial choice and switched fingers, making participants adapt (Potential VP correction). See also Supplementary Video. **B)** In the Spatial task, participants used the analogic stick of the controller to control the trajectory of their right virtual hand (forward, left, and right) to reach and press one of the two buttons. The participants’ virtual hand was programmed to automatically raise the right index when the hand was close to the target button. Participants were asked to reach and press either the button on the same side of the VP (Imitative Block) or the one placed in the opposite space (Complementary Block). Along the trial, the VP would initially move towards a target and in 30% of the trials correct its initial trajectory, making participants adapt (Potential VP correction). See also Supplementary Video.

By implementing these tasks, and by locking the EEG signals to either the VP’s corrections and the participants’ behavioral adaptation to the VP’s corrections, we aimed at unravelling self and other-related monitoring activity during motor interactions. More specifically, we tested the hypothesis that during motor interactions the monitoring system dedicates its resources to the interaction success rather than to one’s sub-goal. Furthermore, we hypothesized that the monitoring process in the *Spatial* task would recruit resources associated with “low-level” error detection (i.e., late Pe) while the unexpected goal corrections in the *Goal* task would generate “high-level” error responses (i.e., ERN and early Pe).

## Results

### Behavioral results

The present work used VR to reproduce realistic interpersonal interactions. To estimate the ecological validity of our tasks (i.e., to confirm that participants predicted the VP’s movements and adapted to its actions), we extracted two separate task-specific behavioral measures. Specifically, for the *Goal* task we extracted the Last Pressed Finger (LPF) variable that represents the time between the go-signal and the choice of the finger used to press the virtual button to complete the task. In case participants corrected their behavior (e.g., chose the index and then switched to the middle finger because of a correction from the VP) the value represents the time of the final finger choice. For the *Spatial* task, the kinematics of the participants’ virtual hand movements were extracted on the axis perpendicular to the direction towards the targets. Subjects’ Trajectory Corrections (STC) were computed to determine the timing of participants’ adaptation following VP’s corrections (see Methods section).

Data were analyzed according to the design factors (when applicable): CORRECTION, dividing trials where the VP did (Correction trials) or did not correct its actions during the trial (NoCorrection trials), and INTERACTION TYPE, depending on the block instructions for participants to perform Imitative or Complementary actions with regards of those of the VP. When parametric assumptions were not met, non-parametric tests were performed.

In the *Goal* task, a Friedman ANOVA comparing the four conditions showed a significant difference across conditions in Last Pressed Finger values (χ2(3) = 42.42, *p* < 0.001). LPF values were significantly longer (*p* value corrected for multiple comparisons = 0.05 / 4 = 0.0125) for Correction trials compared to NoCorrection trials both in Imitative (*p* < 0.001) and Complementary trials (*p* < 0.001; other comparisons *p >* 0.31). As no significant difference was detected between Complementary and Imitative trials, in Figure 2A we show the averaged LPF values for Correction (M = 0.715s ± 0.016) and NoCorrection conditions (M = −0.021s ± 0.068). Negative times represent LPF happening before the occurrence of a correction from the partner (or the same time-point in NoCorrection trials).

**Figure 2.**
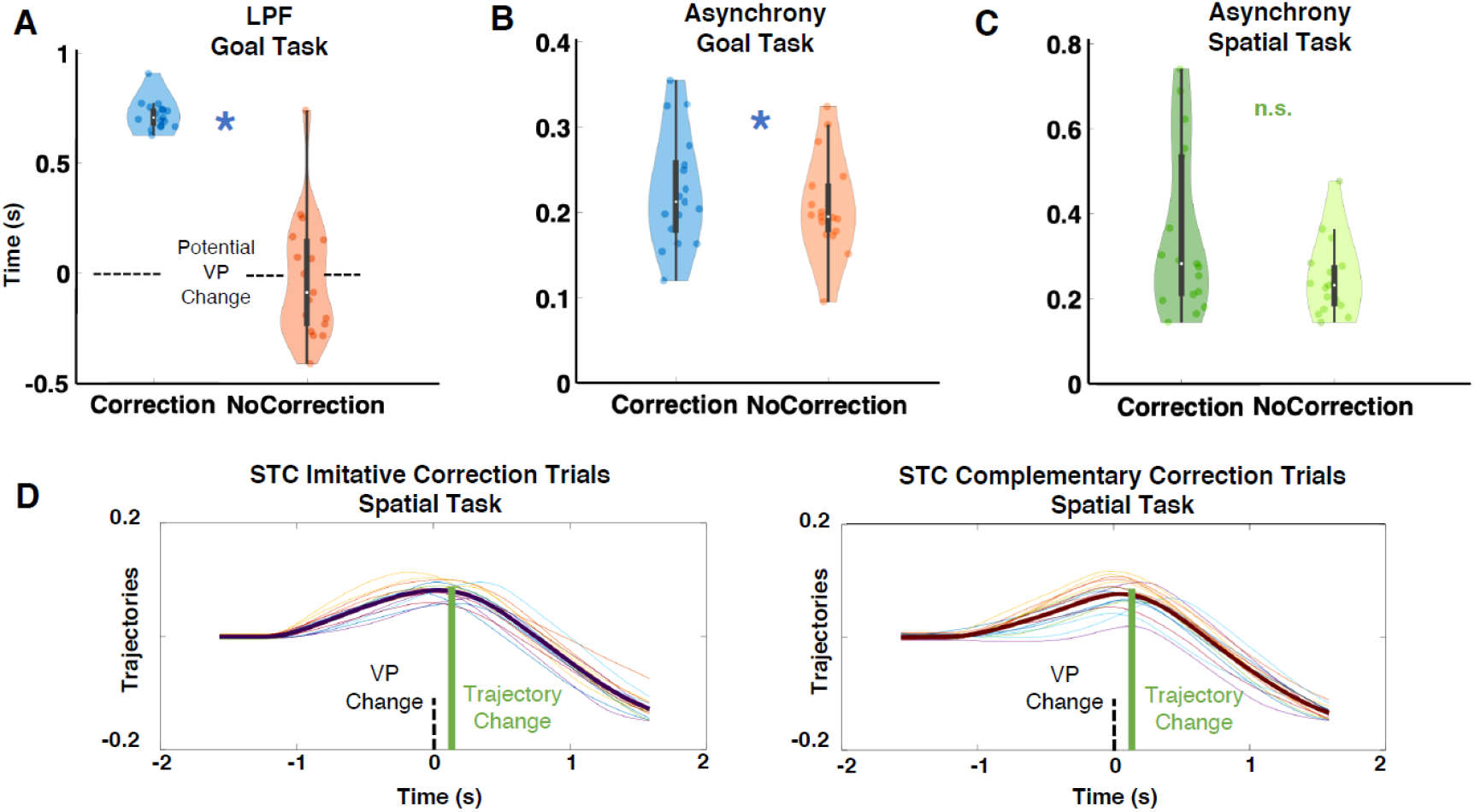
Behavioral results. A) Task-specific behavioral measure for the Goal task. Last Pressed Finger (LPF) values represent the time of the final choice regarding the finger used to press the virtual button. In Correction trials, participants had to correct their behavior following an unexpected goal change from the VP. Time 0 on the Y axis corresponds to the frame where the VP did or did not switch fingers (Potential VP correction). Asynchrony represents the absolute value of the time delay between subjects' and VP’s pressing times in **B)** the Goal task and **C)** the Spatial task. **D)** Subjects trajectories in Correction trials in both Imitative and Complementary conditions. Shaded lines represent single subjects’ trajectories and bold lines represent the across-subjects’ mean trajectory. Time 0 on the X axis corresponds to the frame where the VP corrected its initial trajectory (VP correction). Subjects’ Trajectory Corrections (STC) were computed to determine the timing of subjects’ adaptation following VP’s corrections; see Methods section. Asterisks indicate significant (p <.05) differences. Violin plots display box plots, data density and single subjects’ values (dots).

In the *Spatial* task, no Subjects’ Trajectory Correction (STC) was detected in NoCorrection trials (see Supplementary Figure 1); therefore, data was analyzed using a dependent-sample t-test comparing STC in Imitative and Complementary actions in Correction trials. The dependent-sample t-test revealed that the STC in correction trials between Imitative (M = 0.086s ± 0.056) and Complementary (M = 0.087s ± 0.031) conditions did not differ (*t*(16) = 0.093, *p* = 0.93; see Figure 2C).

These task-specific markers of behavioral adaptation confirm online adjustments to the unexpected movements from the VP. Specifically, in the *Goal* task during Correction trials participants were forced to switch their own action type following VP’s corrections, resulting in longer Last Pressed Finger times compared to NoCorrection trials. Similarly, in the *Spatial* task, kinematics data show a clear trajectory change during Correction trials (and not in the NoCorrection ones, see Supplementary Figure 1), indicating that subjects were mirroring the trajectory of the VP before the correction. Altogether, one can assume that the two tasks simulated a rather realistic environment that led to ecological behaviors from the participants.

In order to quantify the participants’ efficiency in performing the interaction tasks we considered subject-VP asynchrony in pressing the virtual button (i.e., the absolute time difference between VP’s and participants’ pressing time, with lower values highlighting better synchronization between the participants’ avatar and the VP). For other behavioral measures such as Reactions times and Movements times, see Supplementary Materials.

In the *Goal* task, the 2 CORRECTION (Correction, NoCorrection) x 2 INTERACTION TYPE (Imitative, Complementary) repeated-measures ANOVA showed a significant main effect of CORRECTION (*F*(1, 16) = 6.669, *p* = 0.020, ηp^2^ = 0.30), highlighting that synchrony was worse in Correction trials (M = 0.22s ± 0.016) compared to NoCorrection trials (M = 0.20s ± 0.013; see Figure 2B). There was no significant main effect of INTERACTION TYPE (*F*(1, 16) = 2.40, *p* = 0.140, ηp^2^ = 0.13) nor interaction of the two factors (*F*(1, 16) = 0.927, *p* = 0.349, ηp^2^ = 0.054).

In the *Spatial* task, the 2 CORRECTION (Correction, NoCorrection) x 2 INTERACTION TYPE (Imitative, Complementary) repeated-measures ANOVA showed that asynchrony was not influenced by any factor or their interaction (overall M = 0.251s ± 0.088) (see Figure 2C).

### EEG results locked to VP’s behavior – Goal Task

The analyses of the EEG activity locked to the VP’s corrections (or to the same time-point in the NoCorrection trials) focused on previously described error-related neuromarkers in the time domain (i.e., ERN, early and late Pe). For results in the time-frequency domain (i.e., midfrontal theta), see Supplementary Materials (Supplementary Figure 2). As described above, these markers highlight different hierarchical processes, with early and frontal components associated with higher-level errors (ERN, early Pe), and later, more posterior, components associated with lower-level error-processing (late Pe). We found the presence of an ERN (100-220 ms) over FCz in trials where the VP corrected its pressing finger unexpectedly.

The 2 CORRECTION (Correction, NoCorrection) x 2 INTERACTION TYPE (Imitative, Complementary) repeated-measures ANOVA on ERN values showed a significant main effect of CORRECTION (*F*(1, 16) = 40.147, *p* < 0.001, ηp^2^ = 0.71), with larger ERN amplitude in Correction trials (M = −1.237µV ± 0.817) compared to NoCorrection trials (M = −0.047µV ± 0.284, see Figure 3A). There was no significant main effect of INTERACTION TYPE nor interaction of the two factors (*ps* > 0.061).

**Figure 3.**
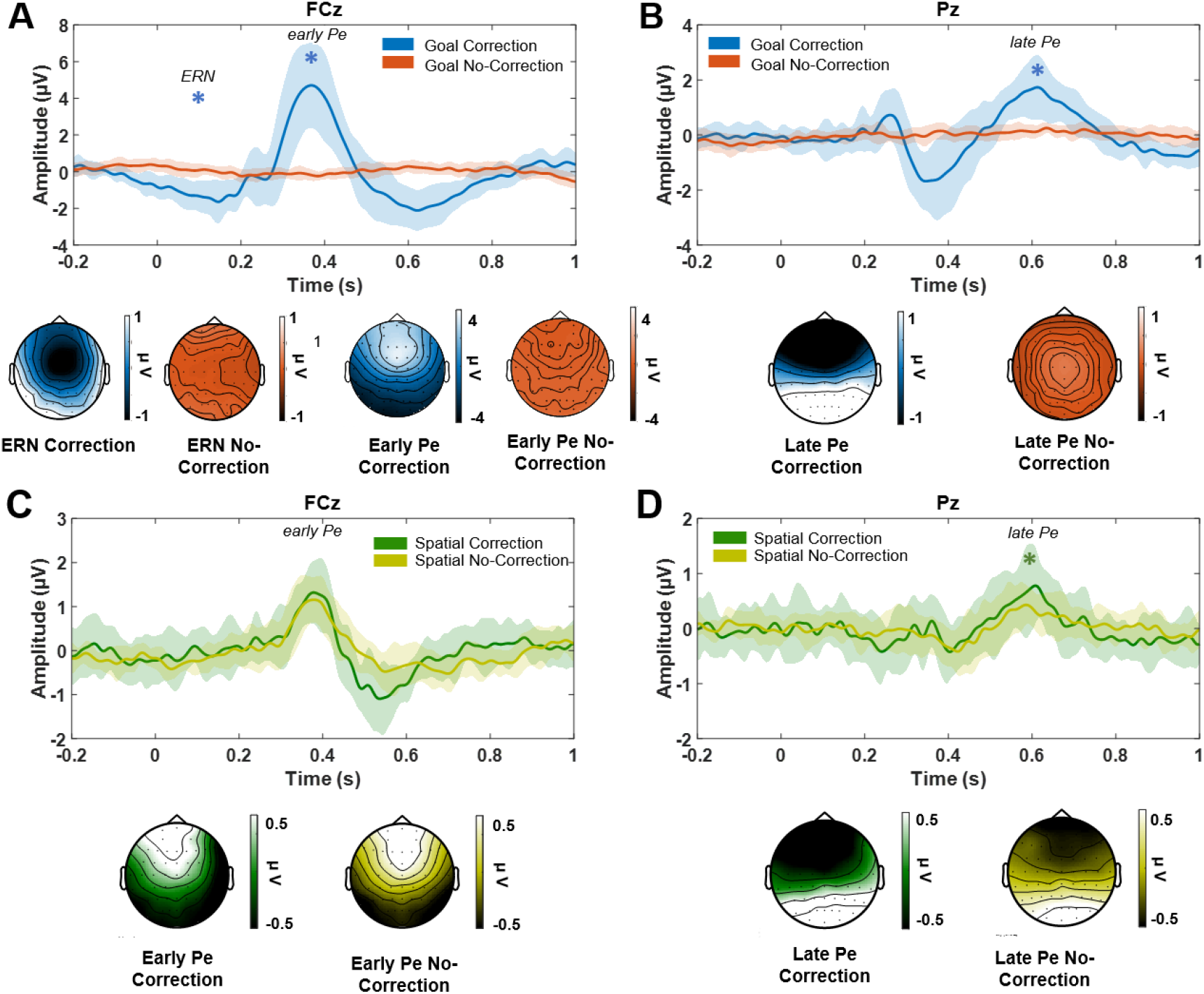
Event-related potentials locked to the VP’s correction or to the same time-points in the absence of corrections in A) the Goal task over FCz, with larger ERN and early Pe amplitudes for Correction vs. NoCorrection trials; **B)** the Goal task over Pz, with larger late Pe amplitudes for Correction vs. NoCorrection trials; **C)** in the Spatial task over FCz, with no difference for early Pe amplitudes between Correction and NoCorrection trials and; **D)** in the Spatial task over Pz, with larger late Pe amplitudes for Correction vs. NoCorrection trials. Asterisks indicate significant (p <.05) differences.

Similarly, the 2 CORRECTION (Correction, NoCorrection) x 2 INTERACTION TYPE (Imitative, Complementary) repeated-measures ANOVA on early Pe (250-450 ms) component over FCz showed a significant main effect of CORRECTION (*F*(1, 16) = 84.997, *p* < 0.001, ηp^2^ = 0.84; other *ps* > 0.148). The early Pe amplitudes were larger in Correction trials (M = 2.747µV ±1.283) compared to NoCorrection trials (M = −0.119µV ± 0.265, see Figure 3A).

The analysis of the late Pe component (500-700 ms) over Pz also showed larger amplitudes for Correction trials (M = 1.228 µV ± 0.698 - see Figure 3B) compared to NoCorrection trials (M = −0.127 µV ± 0.196; *F*(1, 16) = 58.554, *p* < 0.001, ηp^2^ = 0.78). No other comparison reached significance (*ps* > 0.177).

### EEG results locked to VP’s behavior – Spatial Task

Contrary to the *Goal* task, no ERN was detected over FCz for Correction trials in the *Spatial* task. Furthermore, the analysis of the early Pe (300-450 ms) over FCz showed that early Pe amplitudes (overall M = 0.789 µV ± 0.439) were not influenced by any factors (*ps* > 0.238, Figure 3C). However, late Pe amplitudes (450-670 ms) over Pz were higher for Corrections trials (M = 0.376 µV ± 0.393) compared to NoCorrection trials (M = 0.174 µV ± 0.269, *F*(1, 16) = 10.408, *p* = 0.005, ηp^2^ = 0.40 - see Figure 3D – other comparisons were non-significant *ps > 0.06*).

### Self-VP distinction - comparing ERPs locked to VP’s correction and subjects’ adaptation

To unravel self and other-monitoring mechanisms, we further focused on Correction trials of both tasks to establish whether the evoked activity was generated by the correction in the VP’s action or by the fact that this change made the ongoing action of the subject wrong and to be re-planned. To do so, single epochs were redefined and locked to the task-specific motor variable previously extracted to determine the timing in which subjects adapted (namely, Last Pressed Finger for the *Goal* task and the Subjects’ Trajectory Correction for the *Spatial* task). This allowed us to compare the EEG times series when locked to the VP’s behavior (‘VP’) or to the subjects’ behavior (‘Self’). In the *Goal* task, all three ERPs showed higher amplitudes when the signal was locked to the VP’s corrections than when locked to the subjects’ LPF. In details, the 2 IDENTITY (Self, VP) x 2 INTERACTION TYPE (Imitative, Complementary) repeated measure ANOVAs showed a significant main effect of IDENTITY for the ERN (*F*(1, 16) = 30.127, *p* < 0.001, ηp^2^ = 0.65; M_ERN_VP_ = −1.237 µV ± 0.817; M_ERN_LPF_ = 0.083 µV ± 0.544, see Figure 4A), the early Pe (*F*(1, 16) = 85.264, *p* < 0.001, ηp^2^ = 0.84; M_ePE_VP_ = 2.745 µV ± 1.283; M_ePE_LPF_ = −0.044 µV ± 0.288) and the late Pe (*F*(1, 16) = 66.317, *p* < 0.001, ηp^2^ = 0.81; M_lPE_VP_ = 1.228 µV ± 0.698; M_lPE_LPF_ = 0.012 µV ± 0.383), with larger amplitudes for the ERPs locked to the VP’s corrections compared to when locked to the participants’ corrections (see Figure 4B).

**Figure 4.**
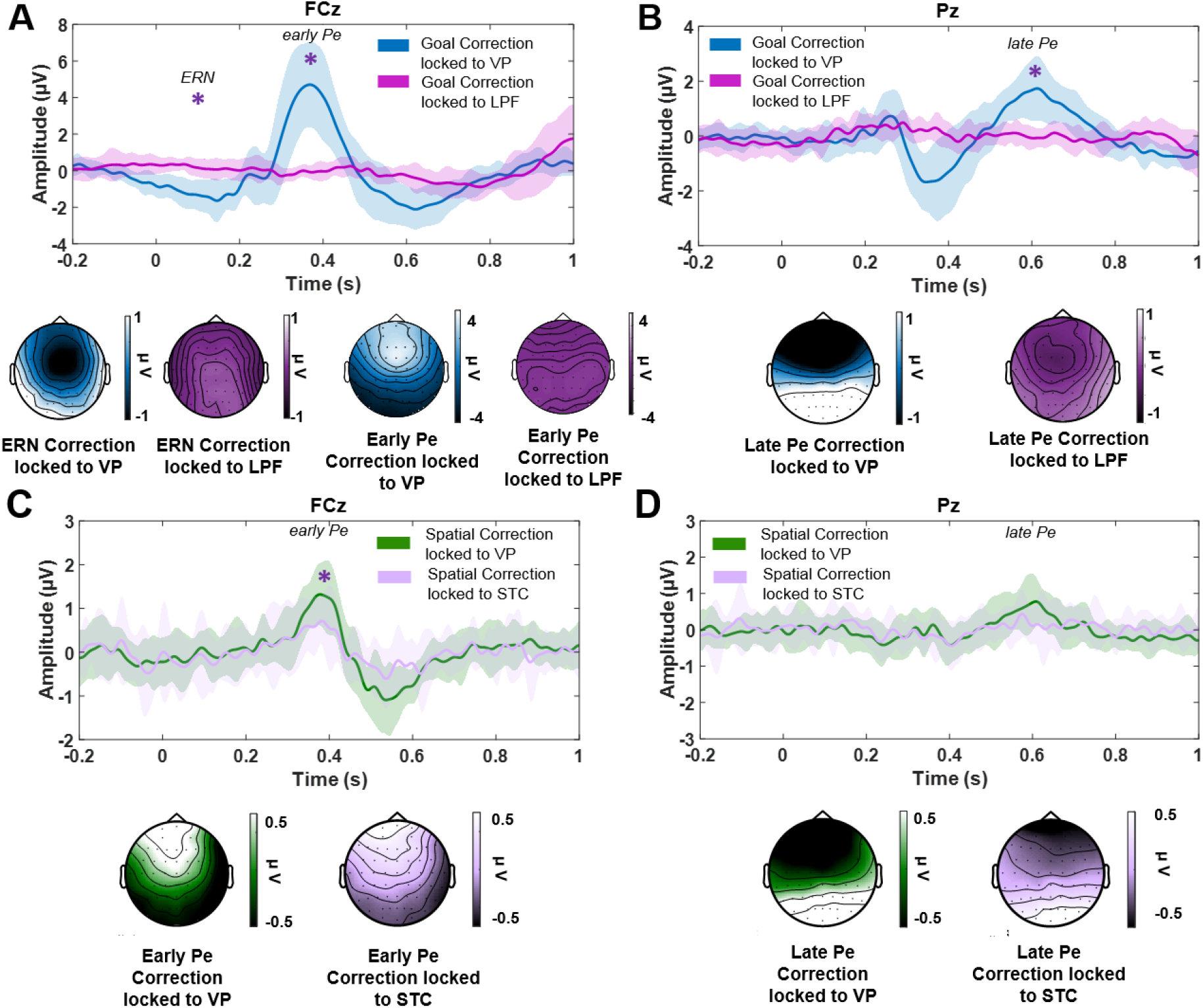
Event-related potentials locked to the VP’s correction (Other) and to the subsequent participants’ adaptation (Self) in A) the Goal task over FCz, with larger ERN and early Pe amplitudes for Correction trials locked to the VP’s corrections vs. locked to the participants’ corrections; B) the Goal task over Pz, with larger late Pe amplitudes for Correction trials locked to the VP’s corrections vs. locked to the participants’ corrections; C) in the Spatial task over FCz, with larger early Pe amplitudes for Correction trials locked to the VP’s corrections vs. locked to the participants’ corrections and; D) in the Spatial task over Pz, no difference in late Pe amplitudes. Asterisks indicate significant (p <.05) differences.

For the *Spatial* task, the 2 IDENTITY (Self, Other) x 2 INTERACTION TYPE (Imitative, Complementary) repeated-measures ANOVA showed a significant main effect of IDENTITY (*F*(1,16) = 7.760, *p* = 0.013, ηp^2^ = 0.33) for early Pe, with larger amplitudes for Correction trials locked to the VP’s corrections (M Me_PE_VP_ = 0.806 µV ± 0.504) compared to Correction trials locked to the subjects’ adaptation (M Me_PE_traj_ = 0.616 µV ± 0.538; see Figure 4C). There was no significant main effect of INTERACTION TYPE nor interaction of the two factors (*ps* < 0.086). For the late Pe, the ANOVA did not show any significant modulation (*ps* < 0.057) (Figure 4D).

## Discussion

In the present study, we recorded EEG in participants that were interacting with a Virtual Partner in fully immersive Virtual Reality scenarios. In two motor interaction tasks involving a synchronous button pressing with a partner, participants controlled the arm and hand movements of a virtual avatar seen from first-person perspective, by means of a joystick. The main advantage of using VR is that participants experienced the feeling of being in control of their virtual body (see data on Ownership and Agency in Supplementary Materials) and being involved in realistic motor interactions while being relatively still. The main goal of these tasks was to target the EEG correlates of monitoring unexpected actions from a partner during social interactions. More precisely, the co-recording of behavioral, kinematics and EEG data allowed us to disentangle neural activity related the self and other-action monitoring. Furthermore, we aimed at distinguishing EEG correlates of “low” and “high-level” motor correction processing during an interaction containing unforeseen actions from a partner, namely a trajectory correction (*Spatial* task, low-level) or a goal change (*Goal* task, high-level), respectively. Our results show that corrections generated larger ERPs when EEG responses were locked to the VP’s correction than when locked to the subjects’ subsequent adaptation. Moreover, “high” and “low-level” behavioral corrections of the partner generated different EEG patterns in individuals that needed to react to these corrections, with the presence of fronto-central activity (ERN, early Pe) during goal change, while no such pattern was detected in the *Spatial* task (only detected in late Pe).

### High and low-level motor corrections during social interactions

Regarding behavioral performance, Synchrony was worse when the VP performed unforeseen actions compared to when no correction was needed in the *Goal* task, but not in the *Spatial* task (although their movement times were longer, see Supplementary Materials). Importantly, our study also highlights the difference in coding *Goal* and *Spatial* corrections at the neural level. *Goal* changes generated a larger ERN over FCz compared to trials where the VP did not correct its goal. The same pattern was observed for early Pe over FCz and late Pe over Pz components (Figure 3A and 3B). A different pattern of results was detected in the *Spatial* task: first, no ERN was clearly identified for spatial corrections and the difference between Correction and NoCorrection trials was only observed for the late Pe over Pz (the presence of an early Pe was detected but no difference was found, see Figure 3C and 3D). Thus, the analysis of behavioral indices and the ERPs suggests a difference in the neural processing of unexpected corrections from the VP depending on the feature impacted by the change. Previous studies showed that movement and goal-related errors differently influence the feeling of controlling one’s own actions (32, 33), here we show that interacting with someone performing those errors differently affects the dyadic performance. Furthermore, earlier research has proposed that the ERN, reflecting activity from the anterior cingulate cortex (ACC; (34, 35)), underlies the detection of “high-level errors” such as the failure to meet a goal (8). Conversely, the (early and late) Pe is thought to originate from a more anterior ACC region and the posterior cingulate–precuneus (36, 37) or prefrontal and parietal sources (11, 29), and is regarded as reflecting the detection of “low-level motor errors” such as action conflict monitoring, including movement correction (4, 5, 9, 38), but also error categorization and awareness (7, 39). Here, we aimed at distinguishing these two types of errors within the context of interpersonal motor interactions. As hypothesized, a correction in the initial action goal from the VP generated an ERN, suggesting that these corrections are coded as “high-level” prediction errors, while an unexpected correction in the initial trajectory (i.e., low-level) is reflected in a later and more parietal component similar to those associated with “low-level” motor errors.

### Self- and Other- related action monitoring during synchronous interactions

Studying unexpected events in synchronous interactive tasks allows one to describe how the action monitoring system dedicates its resources to the processing of self- and others’ actions. This type of experiments represents a technical challenge (40–42) and has important theoretical and practical repercussions, as they are informative on real-life situations that healthy individuals and patients face in everyday social situations such as sports, musical performance or rehabilitation routines (43, 44). At the theoretical level, defining an error (and the related neural responses to it) at the dyad level is complex as it needs to encompass the evaluation of the interaction (here performing complementary or imitative movements) as well as the monitoring of one's own and other's actions (sub-goals (45)). In this sense, the correction observed in the action of the other can be conceptualized as “interactive”, since its occurrence immediately makes the action of the participants wrong and requires adjustments to achieve both their individual sub-goal (i.e., perform the individual’s correct action) and the dyad goal (i.e., realize a complementary/imitative interaction in synchrony).

The overlap between monitoring self and others’ actions has been studied in the framework of music-making (46) or in sequential inter-agent coordination with an emphasis on the role of anticipatory and adaptation mechanisms (47). Very recently (30), it was shown that hearing and performing various types of music errors (i.e., wrong notes or extra, unexpected notes) while playing with a partner generates different EEG responses, no matter whether the error was unambiguously attributed to one of the two partners or ambiguously perceived as generated by oneself or a partner.

Here, to study the neural dynamics associated with unexpected events during a joint performance, we aimed at disentangling the neural processes related to self- and other-monitoring. Thanks to the use of Virtual Reality and the continuous recordings of high temporal resolution EEG and motion kinematics data, we managed to extract the neural activity locked to either the correction of the VP’s action or to the consequent behavioral adaptation from the participants (i.e., Last Pressed Finger in the *Goal* task or Subjects’ Trajectory Corrections in the *Spatial* task). Overall, the ERPs associated with error-processing were larger when neural responses were locked to the VP’s correction than when locked to the subjects’ adaptation (see Figure 3 and 4). In detail, goal changes induced larger ERN, early and late Pe when locked to the partner’s correction rather than to the self-adaptation; spatial corrections induced a larger early Pe when locked to the partner’s correction compared to the self-correction. Therefore, we show that correction-related EEG markers are likely due to the violation of the others’ action prediction rather than an internal error signal linked to the re-planning of one’s own action. These results emphasize a larger other-rather than self-monitoring during synchronous motor interactions and highlight a social mode adopted by the monitoring system to handle motor interactions.

## Limitations

During the last decades, many studies have been using Virtual Reality as a tool to study human behavior and neural dynamics as it allows greater experimental control and ecological validity, especially in the case of social interactions (48). For example, mutual adaptation between VPs and human participants were made possible (41, 49–51). The tasks used in the present study allowed us to selectively study the behavioral and electrophysiological markers associated with follower-role taking during motor interactions. Indeed, the participants were not involved in an action-perception closed-loop (27) and were dependent on the VP’s movements. Hence the tasks used in the present study assigned a “leader” role to the VP and a “follower” role to the participants, a condition that might influence behavioral and neural patterns (52–54). Thus, our results may shed light only on follower-related monitoring processes. Future studies should aim at distinguishing processes involved in other dyadic synchronization strategies, such as mutual adaptation (55, 56) or leading–leading interactions (57).

## Conclusion

The present study offers new insights on monitoring dynamics in motor interactions, showing that observing a partner violating action goal prediction elicits frontal and earlier neural responses compared to those associated to others’ spatial corrections. We also show a prominent role of other-rather than self-monitoring mechanisms during interpersonal motor interactions. This evidence not only confirms a hierarchy among the features that need to be monitored during an interaction with a partner but also indicates that the action monitoring system is mainly dedicated to signaling the partner’s behavioral correction rather than the subsequent adaptive correction of one’s own behavior. Finally, the present study shows that participants placed in a virtual environment generate similar brain patterns to the ones observed in classic Human-VP set-ups (58).

## Materials and Methods

### Participants

Eighteen participants (11 females, age: 26.2 years old) took part in the experiment. One subject was removed from the analysis due to technical problems. The final sample comprises seventeen participants (10 females, age: 26.8 years old). The sample size was selected by means of a power analysis performed with the software More Power 6.0.4. The partial eta squared obtained in a previous study investigating the electroencephalographic markers of error monitoring during motor interactions (0.33) was used as excepted effect size (58). The analysis indicated that a partial eta squared of 0.33, a power of 0.80, an alpha of 0.05 in a 2×2 design required a sample of 18 participants. All participants were right-handed with normal or corrected-to-normal vision. Participants were naïve as to the aim of the experiment at the outset and were informed of the purpose of the study only after all the experimental procedures were completed. The experimental procedures were approved by the Ethics Committee of the Fondazione Santa Lucia (Rome, Italy) and the study was performed in accordance with the 2013 Declaration of Helsinki. All participants gave written consent prior to the experiment.

### Experimental stimuli

The virtual scenario and bodies were designed using 3DS Max 2017 (Autodesk, Inc.) and IClone 7 (Reallusion, Inc.), and implemented in the Unity 5 game software environment. The scenario was presented by means of the Oculus Rift Head Mounted Display (HMD; https://www.occulus.com) having 110° field-of-view (diagonal FOV) with a resolution of 2160 x 1200. The virtual scene consisted of a real-size room (1:1 scale), two virtual avatars sitting on opposite sides of a table and a virtual grey panel placed between the avatars so that the faces were not visible. In front of both avatars, at the center of the table, appeared the 3D model of two buttons (either two identical red rectangles in the *Spatial* task or a yellow cylindric and a purple round button in the *Goal* task), placed on the left and right of the avatar’s body midline, and a LED light that could either turn red or green to signal un successful or successful outcome, respectively (see Figure 1, as well as the Supplementary video). Participants observed the gender-matched virtual body from a 1^st^-person perspective (1PP) through HMD. A right Oculus Touch controller (https://www.oculus.com) was used in order to allow the participants to control the movement of the right arm of their avatar in real time, observed in 1PP. Participants could move the avatar’s hand by using the analogic stick of the Oculus touch controller with their own right thumb in all the directions in the *Spatial* task (i.e., multidirectional control), and only forward in the *Goal* task (i.e., unidirectional control; see Figure 1). During the experiment, the virtual scenario was rendered in both HMD and a computer screen, such that the experimenters could observe the participants performance.

### Procedure

All participants were seated comfortably on a chair in front of a table and wore the HMD (through which they observed the gender-matched virtual body and the scenario in 1PP), as well as a 64-channel EEG cap. Before each task, participants underwent Calibration, Familiarization and Training phases. In the Calibration phase, the perspective point-of-view of each participant was adjusted to pair the 1PP “Self-avatar” virtual body with individual positioning in order to obtain the best spatial-match between the participant’s real and virtual body. In the Familiarization phase, participants were invited to look both at their virtual body (“Self-avatar”) and at the environment, and to verbally describe what they were seeing (~30 sec; (59)). In the Training phase, participants were required to control their avatar’s right hand with the Oculus touch controller (see Figures 1A and 1B) to reach and press the button as synchronous as possible with the virtual partner (~ 10 trials). The 3D LED light placed on the table gave visual feedback to the participants about their performance, i.e., if they pushed the button synchronously with the virtual partner the light turned on in green; if they failed, the light turned red. During both the Training phase and the Experimental procedure, the timing determining if the trial was successful was adjusted trial by trial following a staircase procedure. In details, for the first trial of each block, the interaction was considered successful (i.e., the lightbulb turning green) if the participants managed to reach the button within a 200ms difference with the VP. If the participants failed, the time window accepted as successful increased of 50 ms (i.e., 250ms). If participants were successful, the time window accepted as successful was shrunk of 50 ms (i.e., 150ms). The timing of the successful interaction was then corrected using steps of +50 or −50 ms throughout the block.

The total procedure was composed of four separate and counterbalanced blocks across participants, namely two blocks for the *Goal* task (Imitative-Complementary, see below) and two blocks for the *Spatial* task (Imitative-Complementary, see below). Putting the thumb on the analogic stick of the right Oculus controller, participants could spatially control the speed of the virtual hand (only forward). Pressing the index and middle trigger button of the controller, participants could raise either the virtual index associated with the round purple button or the virtual middle finger, associated with the cylindric yellow button on their avatar side (see Figure 1A). In this condition, participants were asked to choose a finger to accomplish the action synchronized with the virtual partner. The *Goal* task was repeated in two conditions requiring participants to reach and press the same-colored button using the same finger as virtual partner (*Goal* task Imitative) or to press a different colored button, using the opposite finger with respect to the virtual partner (*Goal* task Complementary, see Figure 1A). The Complementary and Imitative blocks were counterbalanced across participants. In both cases, the aim of the trial was to push the button as synchronously as possible with the Virtual Partner. Participants were instructed before each block to either imitate or complement the action of the VP. In the *Spatial* task, two red rectangular buttons (a right and a left one), were placed at the midline of the virtual table (see Figure 1B). By using the analogic stick of the right Oculus controller, participants could spatially control the trajectory of their right virtual hand (forward, left and right) and thus reach and press one of the two buttons. The participants’ virtual hand was programmed to automatically raise the right index when the hand was close to the target button. Thus, in this block, participants were required only to use their right thumb to move the right virtual hand and arm and press the button as synchronously as possible with the virtual partner. The *Spatial* task was repeated twice where participants were asked to reach and press the same button in space of the VP (*Spatial* task Imitative) or the one placed in the opposite part (*Spatial* task Complementary, Figure 1B). Participants were instructed before each block to either imitate or complement the action of the VP. Each block was presented in a counterbalanced order across subjects and consisted of 100 trials per block (total 400 trials). Crucially, in 30% of the trials of each block, the Virtual Partner corrected its initial behavior, i.e., from aiming to press the round purple button using the index to pressing the squared yellow button with the middle finger (*Goal* task) or going from right to left (*Spatial* task), therefore forcing the subject to change his/her initial action to perform the task correctly. The EEG was time-locked (time 0) to the Virtual Partner’s correction time (or the corresponding time in no-correction trials) (see Figure 1).

Each trial started with an acoustic ‘go’ signal (“beep”) delivered through the Oculus headphones. Both avatars started with their hands closed and placed in the center of table’s midline. Depending on the block, participants had to either imitate or complement the action of the Virtual Partner either in space (*Spatial* task) or at a goal-level pressing the yellow or purple button using their index or their middle finger (*Goal* task) while being as synchronous as possible in pressing the button they had to reach. Once both the virtual partner and the subject’s avatar pressed their respective buttons, the LED either turned green or red, depending on the absolute time difference between the two pressing times. As said above, the visual feedback was also governed by a plus or minus 50 ms staircase procedure. The trial ended 2 seconds after the LED visual feedback. Crucially, as mentioned before, the Virtual Partner could change its initial behavior (Correction Factor). The timeline of a trial in the *Goal* task was the following: after the go signal (“beep”) was provided, the virtual partner started its movement which lasted 3170 ms. 1056 ms after the beginning of the movement (i.e., at 33% of the whole movement time), the VP raised a finger in order to press the associated button; in the case of a correction trial, the virtual partner changed the initial finger 2113 ms after starting its movement (i.e., at 66% of the whole movement time, see Figure 1). The timeline of a trial in the Spatial task was the following: after the go signal (“beep”) was provided, the virtual partner started its movement to reach the left or the right button (movement time lasting 3170 ms); in the case of a correction trial, the virtual partner changed the trajectory 1585 ms after starting its movement (50% of the whole movement time, see Figure 2).

### Behavioral results

Synchronicity values (an index of performance efficiency corresponding to the absolute time difference between the VP and the subject’s touching time), were computed automatically for each trial.

The task-specific behavioral measures were extracted in order to determine when the subject made their final choice regarding which finger (index or middle finger, *Goal* task) or direction (right or left, *Spatial* task) to use to solve the task.

For the *Goal* task the Last Pressed Finger (LPF) variable represents the time between the go-signal and the choice of the finger used to press the virtual button to complete the task. As explained in the Result section, in case participants corrected their behavior, for example chose the index and then switched to the middle finger (following a VP correction), the value represents when the final finger choice was made, in this case the middle finger. The constant value of 2.113s (time between the go-signal and the crucial frame when the VP could switch its action) was subtracted from the LPF values so that negative values highlight subjects’ decision before the actual frame where the VP corrected its behavior (in correction trials) and positive values reflect an adaptation to the VP’s correction.

For the *Spatial* task, the kinematics of the virtual hand movement on the plane of the movements were extracted, and trajectory profile were computed. To determine the timing when subjects adapted and corrected directions (following VP’s corrections), we performed a peak detection analysis on single correction trials using the Matlab function ‘findpeak.m’.

### EEG pre-processing

EEG signals were recorded and amplified using a Neuroscan SynAmps RT amplifiers system (Compumedics Limited, Melbourne, Australia). These signals were acquired from 60 tin scalp electrodes embedded in a fabric cap (Electro-Cap International, Eaton, OH), arranged according to the 10-10 system. The EEG was recorded from the following channels: Fp1, Fpz, Fp2, AF3, AF4, F7, F5, F3, F1, Fz, F2, F4, F6, F8, FC5, FC3, FC1, FCz, FC2, FC4, FC6, T7, C5, C3, C1, Cz, C2, C4, C6, T8, TP7, CP5, CP3, CP1, CPz, CP2, CP4, CP6, TP8, P7, P5, P3, P1, Pz, P2, P4, P6, P8, PO7, PO3, AF7, POz, AF8, PO4, PO8, O1, Oz, O2, FT7, and FT8. Horizontal electro-oculogram (HEOG) was recorded bipolarly from 2 electrodes placed on the outer canthi of each eye and signals from the left earlobe were also recorded. All electrodes were physically referenced to an electrode placed on the right earlobe and were algebraically re-referenced off-line to the average of all electrodes. Impedance was kept below 5 KΩ for all electrodes for the whole duration of the experiment, amplifier hardware band-pass filter was 0.01 to 200 Hz and sampling rate was 1000 Hz. To remove the blinks and eyes saccades, EEG and horizontal electro-oculogram were processed in two separate steps. Data were then down sampled at 500 Hz before a blind source separation method was applied on continuous raw signal, using Independent Component Analysis (ICA, (60) implemented in the Fieldtrip toolbox to remove any components related to eye movements from the EEG. 59 components were generated and artifactual components (blinks and saccades) were removed based on the topography and the explained variance (components are ordered by the amount of variance they represent), data were then visually inspected after the removal of the components, to check if both blinks and saccades were no longer present (1.63 components per participants were rejected on average).

### ERPs extraction

Time domain analyses were performed by using the FieldTrip (version 2019-01-13) routines (Donders Institute, Nijmegen; (61) in Matlab2017a (The MathWorks, Inc.). For both the *Goal* and *Spatial* tasks, the pre-processed EEG time series were band-pass filtered (0.5 to 40 Hz) and 2s epochs were created in the −1000 ms to 1000 ms period, time zero being the Correction/NoCorrection frame (see Figure 1). Each trial was then baseline corrected from 200 ms to 0 ms before the VP’s correction (or absence of correction). ERPs mean amplitudes for each subject were extracted at FCz for ERN/early Pe, and Pz for late Pe according to previous literature. The time windows were determined based on visual inspections of the grand averages, and consisted in: 1) 100-220 ms over FCz for the ERN, 2) 250-450 ms over FCz for the early Pe and 3) 500-700 ms over Pz for the late Pe in the *Goal* task (see Figure 3A and 3B); 4) 300-450 ms over FCz for the early Pe and 5) 450-670 ms over Pz for the late Pe in the *Spatial* task (see Figure 3C and 3D). The second analysis was performed on Correction trials of both tasks in order to establish whether the evoked activity was generated by the correction of the VP’s action or by the fact that its change made the behavior of the subject wrong and needing adaptation to try and be successful. To do so, single epochs were redefined and locked to the task-specific motor variable previously extracted to determine the timing in which subjects adapted (namely, Last Pressed Finger for the Goal task and the Trajectory change for the Spatial task). This allowed us to compare the EEG times series when locked to the VP’s behavior or to the subjects’ behavior (see Figures 4).

### Statistics and Data handling

After EEG artifact rejections, the removal of trials where subjects did not perform accurately in regards to the task (e.g. performing an imitative action when asked to perform a complementary one) or with behavioral outlier detections (i.e., behavioral value that fell above or below 2.5 SDs each individual mean for each condition), the remaining average number of trials for each condition and block is as follows: 77.18% (± 13.32%) trials for Spatial-Imitative, 78.24% (± 8.59%) trials for Spatial-Complementary, 86.29% (± 9.29%) trials for Goal-Imitative and 87.00% (± 7.33%) trials for Goal-Complementary block. Because of violations of ANOVA assumptions, Last Pressed Finger values were analyzed through a non-parametric Friedman ANOVA and follow-up comparisons with Wilcoxon paired tests. Furthermore, we found that no trajectory correction was detectable in NoCorrection trials for the *Spatial* task. Hence, this variable was analyzed by means of a dependent-sample t-test comparing Imitative and Complementary trials only. The rest of the behavioral variables and ERPs’ amplitudes locked to the VP’s behavior met the assumptions for parametric testing and were analyzed separately for the Goal and the Spatial tasks through a 2 × 2 repeated measures ANOVA, with CORRECTION (Correction, NoCorrection) and INTERACTION TYPE (Imitative, Complementary) as factors. The comparison between ERPs locked to the VP’s correction and the subjects’ correction for Correction trials was assessed through a 2 } 2 repeated measures ANOVA, with IDENTITY (Self, VP) and INTERACTION TYPE (Imitative, Complementary) as factors. All statistical tests were carried out in Statistica (StatSoftware).

## Acknowledgments

The authors thank Dr. Duru Gün Özkan for her support in the earlier stages of this research. The authors also thank Mr. Ronan McCann for proofreading the manuscript.

## Supplementary Materials

### Behavioural Measures

Several extra behavioral measures were considered. Reaction times (RTs) correspond to the time between the starting “beep” and the subject’s initiating the movement of his/her virtual arm using the joystick. Since subjects could not know whether the VP would correct or not its action at this time of the trial, Correction and NoCorrection trials were collapsed.

In the Goal task, the dependant-sample t-test revealed that the reaction times between Imitative (M = 0.445s, SD = 0.113) and Complementary (M = 0.415s, SD = 0.158) conditions did not differ (t(16) = 0.643, p = 0.52). In the Spatial task, the dependant-sample t-test revealed that the reaction times between Imitative (M = 0.475s, SD = 0.146) and Complementary (M = 0.474s, SD = 0.143) conditions did not differ (t(16) = 0.025, p = 0.98).

Movement Times (MTs) correspond to the time between the starting “beep” and the subject’s touching time. In the Goal task, the 2 CORRECTION (Correction, NoCorrection) x 2 INTERACTION TYPE (Imitative, Complementary) repeated-measures ANOVA showed a main effect of CORRECTION (F(1, 16) = 19.598, p < 0.001, ηp2 = 0.55). Movement times were longer in Correction trials (M = 2.816s, SD = 0.036) compared to NoCorrection trials (M = 2.763s, SD = 0.035). There was no significant main effect of INTERACTION TYPE (F(1, 16) = 0.29, p = 0.867, ηp2 = 0.001) nor interaction of the two factors (F(1, 16) = 0.417, p = 0.527, ηp2 = 0.025).

In the Spatial task, The 2 CORRECTION (Correction, NoCorrection) x 2 INTERACTION TYPE (Imitative, Complementary) repeated-measures ANOVA showed a main effect of Correction (F(1,16) = 12.161, p = 0.003, ηp2 = 0.43). Movement times were longer in Correction trials (M = 2.789s, SD = 0.044) compared to NoCorrection trials (M = 2.625s, SD = 0.051). There was no main effect of INTERACTION TYPE (F(1, 16) = 0.562, p = 0.464, ηp2 = 0.033) nor interaction of the two factors (F(1, 16) = 0.239, p = 0.632, ηp2 = 0.014).

### Subjects’ Trajectory Correction – Spatial task

For the *Spatial* task, the kinematics of the participants’ virtual hand movements were extracted on the axis perpendicular to the direction towards the targets.

**Supplementary Figure 1.**
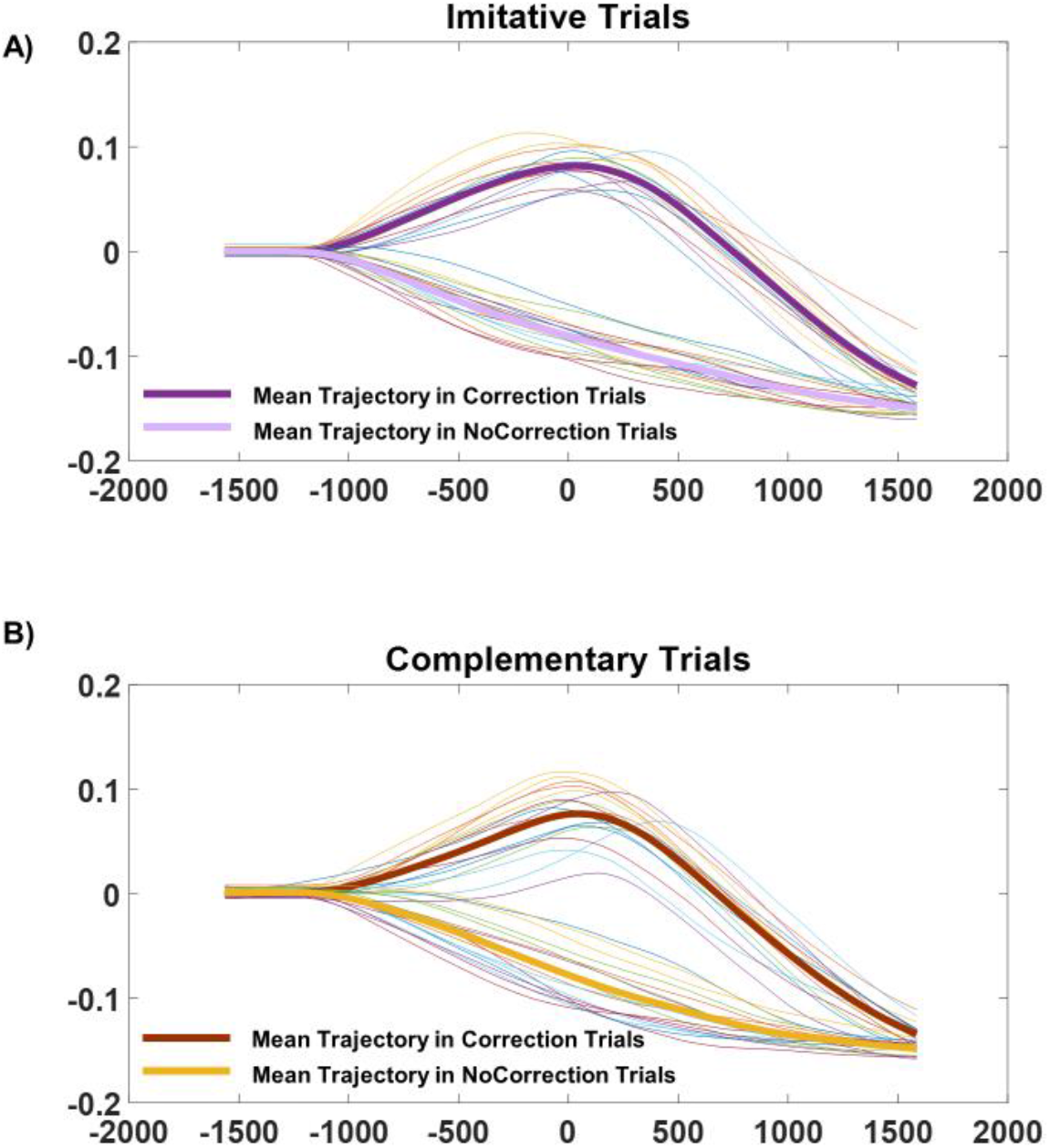
Spatial task trajectories in **A)** Imitative and **B)** Complementary trials. As said in the main text, no trajectory correction was detected in NoCorrection trials.

### Embodiment Ratings

After each task, a black panel with a horizontal green line (60 cm length, left and right extremity marked as “0” and “100” respectively) was presented in the virtual scenario. To assess the degree to which participants experienced the illusory Feeling of Ownership (FO) and Agency (A) over the virtual right hand, a 6-item questionnaire adapted from previous studies (59) was used. The questionnaire consisted of two blocks, each with two items concerning FO (“*I felt as if I were looking at my own hand*”, “*I felt as if the Virtual hand was my hand*”) and a control a FO question (“*It felt as if I had more than one right hand*”); and Agency (“*It felt as if the movements of the Virtual hand were my own movements*”, “*I felt as if I could have caused a/the movement of the Virtual hand*”) and an Agency control question (“*I felt as if the Virtual hand was controlling me*”). Participants were asked to move a vertical bar along the horizontal VAS line by using the analogic stick of the right Oculus touch controller in order to answer the items.

*Goal task - Ownership.* Friedman ANOVA showed a significant effect in Ownership ratings (_X_2(3) = 38.92, *p* < 0.001). As expected, Ownership was higher for Feeling of Ownership questions compared to the control ones (*ps* < 0.001).

*Goal task - Agency.* Friedman ANOVA showed a significant effect in Agency ratings (_X_2(3) = 39.45, *p* < 0.001). As expected, Agency was higher for Agency questions compared to the control ones (*ps* < 0.001).

*Spatial task - Ownership*. Friedman ANOVA showed a significant effect in Ownership ratings (_X_2(3) = 40.11, *p* < 0.001). As expected, Ownership was higher for experimental questions compared to the controls (*ps* < 0.001).

*Spatial task - Agency*. Friedman ANOVA showed a significant effect in Agency ratings (_X_2(3) = 43.92, *p* < 0.001). As expected, Agency was higher for experimental questions compared to the controls (*ps* < 0.001).

### Time-Frequency Analysis

Time-Frequency analyses were performed by using the FieldTrip routines (Donders Institute, Nijmegen; Oostenveld et al., 2010) in Matlab2017a (The MathWorks, Inc.). The EEG time series were obtained by segmenting the signal into epochs of 2500 ms length (from 1000 ms before to 1500 ms after the Correction of the VP). Each epoch was transformed in the frequency domain and multiplied with the Fast Fourier Transformation (FFT) power spectrum of a set of complex Morlet wavelets using 50 ms time bins in a −1000 to 1500 ms peri-stimulus window. Prior to the analysis, data were baselined using a time window from −500 to 0 ms and using the absolute baseline function implemented in Fieldtrip (cfg.baselinetype = ‘absolute’). We conducted whole-scalp analysis using cluster-based nonparametric permutation tests as implemented in the Fieldtrip toolbox. Frequency power in the theta band (4-7 Hz) was averaged and we analysed the change of power for each time bin of 50 ms from 0 to 1000 ms. With the use of dependent samples t-test, clusters of electrodes were generated when *t* values above the significant threshold in at least two of the neighbouring channels were observed. 1000 permutations were run to assess the significance of clusters using a Monte Carlo estimation of significance (*P* < 0.025). Trials were averaged to directly test the comparison Correction vs NoCorrection in both the *Goal* and the *Spatial* tasks.

Since previous studies (53) showed that error-related EEG markers were not affected by the Imitative/Complementary comparisons, these levels were collapsed to increase the signal-to-noise ratio in our time-frequency analysis in the theta band (4-7 Hz). Hence, the cluster-based permutation analysis tested the comparison between Correction and NoCorrection trials and revealed a significant cluster of electrodes (*p* < 0.001) showing higher theta power for Correction trials between 100 and 700 ms after VP correction in the *Goal* task. In line with previous research, the cluster initially showed a fronto-central topography (FCz, FC1, Cz) before expanding to the most part of the scalp (Supplementary Figure 2A). The comparison between Correction and NoCorrection trials in the *Spatial* task revealed a significant cluster of electrodes (*p* < 0.001) showing lower theta power for Correction trials between 350 and 1000 ms after VP correction (Imitative and Complementary conditions collapsed). The cluster initially showed a left occipito-parietal and right fronto-parietal topography before expanding to the most part of the scalp (Supplementary Figure 2B).

**Supplementary Figure 2.**
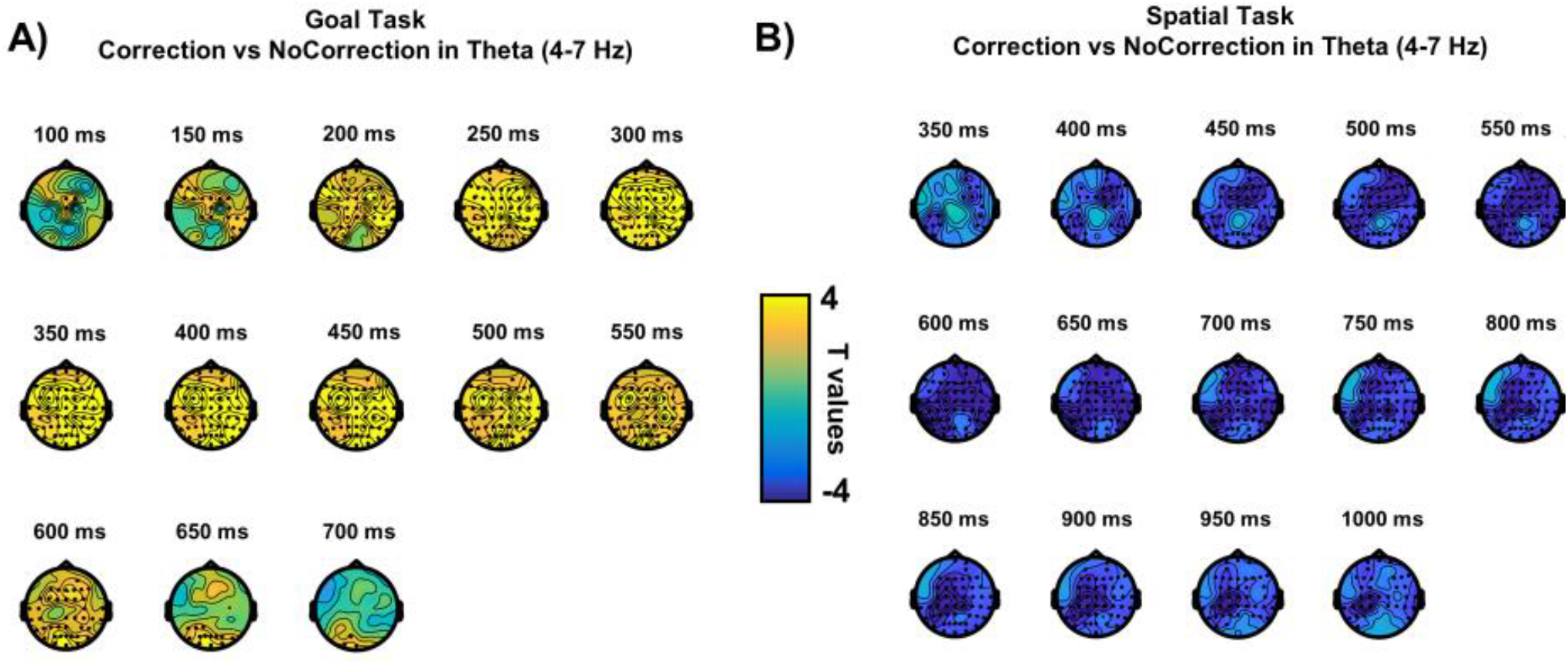
Cluster-based permutation test on theta power between Correction and NoCorrection trials in the **A)** Goal task and **B)** the Spatial task. Dots represent electrodes belonging to significant cluster (p < 0.05) across time highlighting a larger theta power for Correction trials compared to NoCorrection ones in the Goal task, and a reverse pattern in the Spatial task.

The analysis of the EEG data in the time-frequency domain provides further information regarding the different brain processes involved in monitoring other’s actions in interactive scenario. The cluster-based analysis revealed a higher theta power (4-7 Hz) for Correction compared to NoCorrection trials in the *Goal* task. The significant cluster seems to differ over fronto-central areas soon after the perception of the partner change before spreading over parietal and occipital sites. This result is in line with numerous studies associating an increase of theta power to error-processing. On the other hand, the results in the theta band for the *Spatial* task reveal a different pattern, with lower theta power for Correction trials compared to NoCorrection ones, and the activity starting to differ over left occipito-temporal and right-fronto-parietal electrodes. The direction of the results in the theta band for the *Spatial* task was not expected. A cautious observation regarding this pattern is that theta power is differently sensitive to goal and spatial changes.

These results expand the current understanding of error monitoring in social interactions. In a previous study, increase in frontal and occipito-temporal theta activity was shown during motor interactions requiring the monitoring of the interactor’s motor behaviour (53). However, in Moreau et al., (53), the goal and direction of the partner’s movement were linked to one another. Here, by dissociating the contribution of goal and trajectory predictions, we can claim that the increase in frontal theta power is specifically related to violations of higher-level goal predictions.

### Supplementary Video

https://www.youtube.com/watch?v=0V2Cam42ivo

## References

1. C. B. Holroyd, M. G. H. Coles, The neural basis of human error processing: Reinforcement learning, dopamine, and the error-related negativity. Psychological Review 109, 679–709 (2002).

2. V. van Veen, C. Carter S., Conflict and Cognitive Control in the Brain (2006) (February 22, 2021).

3. N. Yeung, M. M. Botvinick, J. D. Cohen, The neural basis of error detection: conflict monitoring and the error-related negativity. Psychol Rev 111, 931–959 (2004).

4. M. Falkenstein, J. Hoormann, S. Christ, J. Hohnsbein, ERP components on reaction errors and their functional significance: a tutorial. Biol Psychol 51, 87–107 (2000).

5. S. Nieuwenhuis, K. R. Ridderinkhof, J. Blom, G. P. Band, A. Kok, Error-related brain potentials are differentially related to awareness of response errors: evidence from an antisaccade task. Psychophysiology 38, 752–760 (2001).

6. R. Verleger, P3b: Towards some decision about memory. Clinical Neurophysiology 119, 968–970 (2008).

7. T. J. M. Overbeek, S. Nieuwenhuis, K. R. Ridderinkhof, Dissociable components of error processing: On the functional significance of the Pe vis-à-vis the ERN/Ne. Journal of Psychophysiology 19, 319–329 (2005).

8. O. E. Krigolson, C. B. Holroyd, Evidence for hierarchical error processing in the human brain. Neuroscience 137, 13–17 (2006).

9. O. E. Krigolson, C. B. Holroyd, Hierarchical error processing: different errors, different systems. Brain Res 1155, 70–80 (2007).

10. P. Luu, D. M. Tucker, S. Makeig, Frontal midline theta and the error-related negativity: neurophysiological mechanisms of action regulation. Clin Neurophysiol 115, 1821–1835 (2004).

11. J. F. Cavanagh, M. X. Cohen, J. J. B. Allen, Prelude to and Resolution of an Error: EEG Phase Synchrony Reveals Cognitive Control Dynamics during Action Monitoring. J. Neurosci. 29, 98–105 (2009).

12. M. X. Cohen, Error-related medial frontal theta activity predicts cingulate-related structural connectivity. NeuroImage 55, 1373–1383 (2011).

13. L. T. Trujillo, J. J. B. Allen, Theta EEG dynamics of the error-related negativity. Clin Neurophysiol 118, 645–668 (2007).

14. J. F. Cavanagh, M. J. Frank, Frontal theta as a mechanism for cognitive control. Trends Cogn Sci 18, 414–421 (2014).

15. G. Fusco, et al., Midfrontal theta transcranial alternating current stimulation modulates behavioural adjustment after error execution. Eur J Neurosci 48, 3159–3170 (2018).

16. H. T. van Schie, R. B. Mars, M. G. H. Coles, H. Bekkering, Modulation of activity in medial frontal and motor cortices during error observation. Nat Neurosci 7, 549–554 (2004).

17. W. Miltner, J. Brauer, H. Hecht, R. Trippe, M. Coles, Parallel brain activity for self-generated and observed errors. undefined (2004) (February 22, 2021).

18. T. Koelewijn, H. T. van Schie, H. Bekkering, R. Oostenveld, O. Jensen, Motor-cortical beta oscillations are modulated by correctness of observed action. Neuroimage 40, 767–775 (2008).

19. E. R. A. de Bruijn, R. I. Schubotz, M. Ullsperger, An event-related potential study on the observation of erroneous everyday actions. Cognitive, Affective, & Behavioral Neuroscience 7, 278–285 (2007).

20. M. S. Panasiti, E. F. Pavone, S. M. Aglioti, Electrocortical signatures of detecting errors in the actions of others: An EEG study in pianists, non-pianist musicians and musically naïve people. Neuroscience 318, 104–113 (2016).

21. R. Pezzetta, V. Nicolardi, E. Tidoni, S. M. Aglioti, Error, rather than its probability, elicits specific electrocortical signatures: a combined EEG-immersive virtual reality study of action observation. Journal of Neurophysiology 120, 1107–1118 (2018).

22. G. Spinelli, G. Tieri, E. F. Pavone, S. M. Aglioti, Wronger than wrong: Graded mapping of the errors of an avatar in the performance monitoring system of the onlooker. NeuroImage 167, 1–10 (2018).

23. S. M. Aglioti, P. Cesari, M. Romani, C. Urgesi, Action anticipation and motor resonance in elite basketball players. Nat Neurosci 11, 1109–1116 (2008).

24. M. Candidi, C. M. Vicario, A. M. Abreu, S. M. Aglioti, Competing Mechanisms for Mapping Action-Related Categorical Knowledge and Observed Actions. Cerebral Cortex 20, 2832–2841 (2010).

25. E. Tomeo, P. Cesari, S. M. Aglioti, C. Urgesi, Fooling the kickers but not the goalkeepers: behavioral and neurophysiological correlates of fake action detection in soccer. Cereb Cortex 23, 2765–2778 (2013).

26. P. M. Hilt, et al., Motor Recruitment during Action Observation: Effect of Interindividual Differences in Action Strategy. Cerebral Cortex 30, 3910–3920 (2020).

27. R. Hari, M. V. Kujala, Brain basis of human social interaction: from concepts to brain imaging. Physiol Rev 89, 453–479 (2009).

28. N. Sebanz, H. Bekkering, G. Knoblich, Joint action: bodies and minds moving together. Trends Cogn Sci 10, 70–76 (2006).

29. B. Tunçgenç, E. Travers, M. T. Fairhurst, Leadership and tempo perturbation affect coordination in medium-sized groups. Scientific Reports 11, 4940 (2021).

30. A. Paas, G. Novembre, C. Lappe, P. E. Keller, Not all errors are alike: modulation of error-related neural responses in musical joint action. Social Cognitive and Affective Neuroscience (2021) https://doi.org/10.1093/scan/nsab019 (February 22, 2021).

31. L. M. Sacheli, M. A. Musco, E. Zazzera, E. Paulesu, Mechanisms for mutual support in motor interactions. Scientific Reports 11, 3060 (2021).

32. R. Villa, E. Tidoni, G. Porciello, S. M. Aglioti, Violation of expectations about movement and goal achievement leads to Sense of Agency reduction. Exp Brain Res 236, 2123–2135 (2018).

33. R. Villa, E. Tidoni, G. Porciello, S. M. Aglioti, Freedom to act enhances the sense of agency, while movement and goal-related prediction errors reduce it. Psychological Research (2020) https://doi.org/10.1007/s00426-020-01319-y (February 22, 2021).

34. C. S. Carter, et al., Anterior cingulate cortex, error detection, and the online monitoring of performance. Science 280, 747–749 (1998).

35. V. van Veen, J. D. Cohen, M. M. Botvinick, V. A. Stenger, C. S. Carter, Anterior cingulate cortex, conflict monitoring, and levels of processing. Neuroimage 14, 1302–1308 (2001).

36. R. G. O’Connell, et al., The role of cingulate cortex in the detection of errors with and without awareness: a high-density electrical mapping study. European Journal of Neuroscience 25, 2571–2579 (2007).

37. C. D. Ladouceur, R. E. Dahl, C. S. Carter, Development of action monitoring through adolescence into adulthood: ERP and source localization. Developmental Science 10, 874–891 (2007).

38. V. van Veen, C. S. Carter, The anterior cingulate as a conflict monitor: fMRI and ERP studies. Physiol Behav 77, 477–482 (2002).

39. T. Endrass, C. Franke, N. Kathmann, Error awareness in a saccade countermanding task. Journal of Psychophysiology 19, 275–280 (2005).

40. L. Schilbach, et al., Toward a second-person neuroscience. Behav Brain Sci 36, 393–414 (2013).

41. G. Dumas, Q. Moreau, E. Tognoli, J. a. S. Kelso, The Human Dynamic Clamp Reveals the Fronto-Parietal Network Linking Real-Time Social Coordination and Cognition. Cereb Cortex 30, 3271–3285 (2020).

42. G. Dumas, Towards a two-body neuroscience. Communicative & Integrative Biology 4, 349–352 (2011).

43. R. Pezzetta, M. Wokke, S. M. Aglioti, R. Ridderinkhof, Doing it Wrong: A Systematic Review on Electrocortical and Behavioral Correlates of Error Monitoring in Patients with Neurological Disorders. Neuroscience (2021) https://doi.org/10.1016/j.neuroscience.2021.01.027 (February 22, 2021).

44. D. G. Özkan, R. Pezzetta, Q. Moreau, A. M. Abreu, S. M. Aglioti, Predicting the fate of basketball throws: an EEG study on expert action prediction in wheelchair basketball players. Exp Brain Res 237, 3363–3373 (2019).

45. L. M. Sacheli, S. M. Aglioti, M. Candidi, Social cues to joint actions: the role of shared goals. Front. Psychol. 6(2015).

46. P. E. Keller, G. Novembre, J. Loehr, “Musical ensemble performance: Representing self, other and joint action outcomes” in Shared Representations: Sensorimotor Foundations of Social Life, Cambridge social neuroscience., (Cambridge University Press, 2016), pp. 280–310.

47. B. Harry, P. Keller, Tutorial and simulations with ADAM: an adaptation and anticipation model of sensorimotor synchronization. Biological Cybernetics 113(2019).

48. X. Pan, A. F. de C. Hamilton, Why and how to use virtual reality to study human social interaction: The challenges of exploring a new research landscape. Br J Psychol 109, 395–417 (2018).

49. G. Dumas, G. C. de Guzman, E. Tognoli, J. A. S. Kelso, The human dynamic clamp as a paradigm for social interaction. PNAS 111, E3726–E3734 (2014).

50. V. Era, S. M. Aglioti, C. Mancusi, M. Candidi, Visuo-motor interference with a virtual partner is equally present in cooperative and competitive interactions. Psychological Research 84, 810–822 (2020).

51. V. Era, S. M. Aglioti, M. Candidi, Inhibitory Theta Burst Stimulation Highlights the Role of Left aIPS and Right TPJ during Complementary and Imitative Human–Avatar Interactions in Cooperative and Competitive Scenarios. Cerebral Cortex 30, 1677–1687 (2020).

52. I. Konvalinka, P. Vuust, A. Roepstorff, C. D. Frith, Follow you, Follow me: Continuous Mutual Prediction and Adaptation in Joint Tapping. Quarterly Journal of Experimental Psychology 63, 2220–2230 (2010).

53. I. Konvalinka, et al., Frontal alpha oscillations distinguish leaders from followers: Multivariate decoding of mutually interacting brains. NeuroImage 94, 79–88 (2014).

54. M. Candidi, A. Curioni, F. Donnarumma, L. M. Sacheli, G. Pezzulo, Interactional leader–follower sensorimotor communication strategies during repetitive joint actions. Journal of The Royal Society Interface 12, 20150644 (2015).

55. V. Era, M. Candidi, M. Gandolfo, L. M. Sacheli, S. M. Aglioti, Inhibition of left anterior intraparietal sulcus shows that mutual adjustment marks dyadic joint-actions in humans. Social Cognitive and Affective Neuroscience 13, 492–500 (2018).

56. V. Era, S. Boukarras, M. Candidi, Neural correlates of action monitoring and mutual adaptation during interpersonal motor coordination: Comment on “The body talks: Sensorimotor communication and its brain and kinematic signatures” by G. Pezzulo et al. Phys Life Rev 28, 43–45 (2019).

57. O. A. Heggli, et al., Transient brain networks underlying interpersonal strategies during synchronized action. Social Cognitive and Affective Neuroscience 16, 19–30 (2021).

58. Q. Moreau, M. Candidi, V. Era, G. Tieri, S. M. Aglioti, Midline frontal and occipito-temporal activity during error monitoring in dyadic motor interactions. Cortex 127, 131–149 (2020).

59. G. Tieri, E. Tidoni, E. F. Pavone, S. M. Aglioti, Mere observation of body discontinuity affects perceived ownership and vicarious agency over a virtual hand. Exp Brain Res 233, 1247–1259 (2015).

60. T. P. Jung, et al., Removing electroencephalographic artifacts by blind source separation. Psychophysiology 37, 163–178 (2000).

61. R. Oostenveld, P. Fries, E. Maris, J.-M. Schoffelen, FieldTrip: Open source software for advanced analysis of MEG, EEG, and invasive electrophysiological data. Comput Intell Neurosci 2011, 156869 (2011).

